# Genetic variation and gene expression across multiple tissues and developmental stages in a non-human primate

**DOI:** 10.1101/092874

**Authors:** Anna J. Jasinska, Ivette Zelaya, Susan K. Service, Christine B. Peterson, Rita M. Cantor, Oi-Wa Choi, Joseph DeYoung, Eleazar Eskin, Lynn A. Fairbanks, Scott Fears, Allison E. Furterer, Yu S. Huang, Vasily Ramensky, Christopher A. Schmitt, Hannes Svardal, Matthew J. Jorgensen, Jay R. Kaplan, Diego Villar, Bronwen L. Aken, Paul Flicek, Rishi Nag, Emily S. Wong, John Blangero, Thomas D. Dyer, Marina Bogomolov, Yoav Benjamini, George M. Weinstock, Ken Dewar, Chiara Sabatti, Richard K. Wilson, J. David Jentsch, Wesley Warren, Giovanni Coppola, Roger P. Woods, Nelson B. Freimer

## Abstract

By analyzing multi-tissue gene expression and genome-wide genetic variation data in samples from a vervet monkey pedigree, we generated a transcriptome resource and produced the first catalogue of expression quantitative trait loci (eQTLs) in a non-human primate model. This catalogue contains more genome-wide significant eQTLs, per sample, than comparable human resources, and reveals sex and age-related expression patterns. Findings include a master regulatory locus that likely plays a role in immune function, and a locus regulating hippocampal long non-coding RNAs (lncRNAs), whose expression correlates with hippocampal volume. This resource will facilitate genetic investigation of quantitative traits, including brain and behavioral phenotypes relevant to neuropsychiatric disorders.

---

Efforts to understand how genetic variation contributes to common diseases and quantitative traits increasingly focus on the regulation of gene expression. Most loci identified through genome-wide association studies (GWAS) lie in non-coding portions of the genome^1^, and are enriched for eQTLs; SNPs that regulate transcript levels, primarily those of nearby genes^2^. This observation suggests that eQTL catalogs may signpost specific variants responsible for GWAS signals^3^.

The majority of known human eQTLs have been identified in lymphocytes or lymphoblastoid cell lines obtained from adults^4^. As normal development and function in complex organisms depends on tightly regulated gene expression at specific developmental stages in specific cell types, most existing datasets describing human transcriptome characterization likely miss data relevant to understanding disease^5^. This lack is particularly striking for brain and behavior disorders, given the inaccessibility of the most relevant tissues in living individuals and the enormous modifications that occur in these tissues across development^6^.

The Genotype Tissue Expression (GTEx) project, using samples obtained from several hundred post-mortem donors^7^, has begun to remedy the lack of human data connecting genotypic variation and multi-tissue transcriptome variation. GTEx provides an eQTL catalog, from multiple tissues, that is the most extensive such resource available^7^. However limitations of GTEx, inherent to human research, motivate the generation and investigation of equivalent resources from model organisms. Advantages of model systems include: (1) the feasibility of controlling for inter-individual heterogeneity in environmental exposures and of minimizing the interval between death and tissue preservation; (2) the practicability of obtaining sizable numbers of multi-tissue samples across a full range of developmental stages; and (3) the opportunity to systematically assess phenotypes of interest in individuals carrying particular eQTL variants. Because of the similarities between humans and non-human primate (NHP) species in behavior, neuroanatomy, and brain circuitry^8,9,10^, NHP eQTLs may be particularly valuable for our understanding of neuropsychiatric disorders.

We report here, in 58 Caribbean vervets (*Chlorocebus aethiops sabaeus*) from the Vervet Research Colony (VRC) extended pedigree, the first NHP resource combining genome-wide genotypes^11^, multi-tissue expression data across post-natal development, and quantitative phenotypes relevant to human brain and behavior, in a setting in which key environmental exposures have been carefully controlled (Online Methods). The Caribbean vervets are an Old World monkey population that has expanded dramatically from a founding bottleneck occurring with the introduction of West African vervets to the Caribbean in the 17^th^ Century^10^; it has experienced a drastic reduction in genetic variation and, like recently expanded human population isolates, displays enrichment for numerous potentially deleterious alleles (Ramensky, unpublished data).

Through necropsies performed under uniform conditions, we obtained both brain and peripheral tissue samples from the 58 vervets included in this study, whose genomes were also sequenced^13^. Using these resources we have delineated cross-tissue expression profiles for seven of these tissues, across multiple developmental stages from birth to adulthood. We identified numerous local and distant eQTLs in each tissue, including a master regulatory locus that, via *IFIT1B,* a gene with a hypothesized role in immune function, modulates expression in blood cells of multiple genes on several chromosomes. Additionally, we demonstrated the relevance of vervet tissue-specific eQTLs to higher-order traits, using hippocampus-specific local eQTLs to identify a set of lncRNAs associated with hippocampal volume, a phenotype related to neuropsychiatric disorders^12^.

## Results

We investigated two datasets. Dataset 1, described previously^13^, consists of gene expression levels obtained by hybridizing all available whole blood-derived RNA samples from the VRC pedigree (N=347) to Illumina HumanRef-8 v2 microarrays, which we used because no vervet arrays are available. After filtering out probe sequences not represented in the vervet genome^14^ or containing common vervet SNPs^11^, we estimated expression levels at 6,018 probes, corresponding to 5,586 unique genes (Supplementary Data 1, Supplementary Table 1). Dataset 2 consists of RNA sequencing (RNA-Seq) reads from seven tissues collected under identical conditions from each of 58 sequenced VRC monkeys (representing 10 developmental stages, from birth through adulthood, Online Methods). Five of these tissues play prominent roles in cognitive and behavioral phenotypes ^15-17^: Brodmann area 46 [BA46], a cytoarchitectonically defined region which encompasses most of the dorsolateral prefrontal cortex (DLPFC); hippocampus; caudate nucleus, a component of the dorsal striatum; pituitary gland; and adrenal gland. The other two tissues (cultured skin fibroblasts and whole blood) are relatively accessible, and thus widely used in studies aimed at identifying biomarkers. We assessed expression of an initial set of 33,994 annotated genes. Before analyzing Dataset 2, we minimized spurious signals by excluding genes expressed in fewer than 10% of individuals or at a level lower than one read per tissue. The gene numbers after this exclusion step are listed by tissue and biotype in Supplementary Table 2. A principal components analysis (PCA) of Dataset 2 showed that, overall, expression levels clustered more by tissue than by individual (Supplementary Fig. 1). Most genes were expressed in multiple tissues; 137 genes demonstrated strong expression in only a single tissue (Supplementary Table 3).

### Multi-tissue Expression Data: Variation By Age, Sex, Cellular Composition, and Technical Factors

The availability, in Dataset 2, of multiple samples from both sexes at each age point enabled us to examine developmental trajectories and sex differences in gene expression for each tissue. To maximize our ability to observe patterns, we conducted PCA on the expression of the 1,000 most variable genes, separately by tissue (Fig. 1). Comparison of the ranks of expression of the orthologs of these genes in matched tissues in humans and rhesus macaques yielded Spearman correlations of between ~0.5-0.8 and ~0.3-0.4, respectively (Supplementary Material and Supplementary Tables 4-6).

**Fig. 1.**
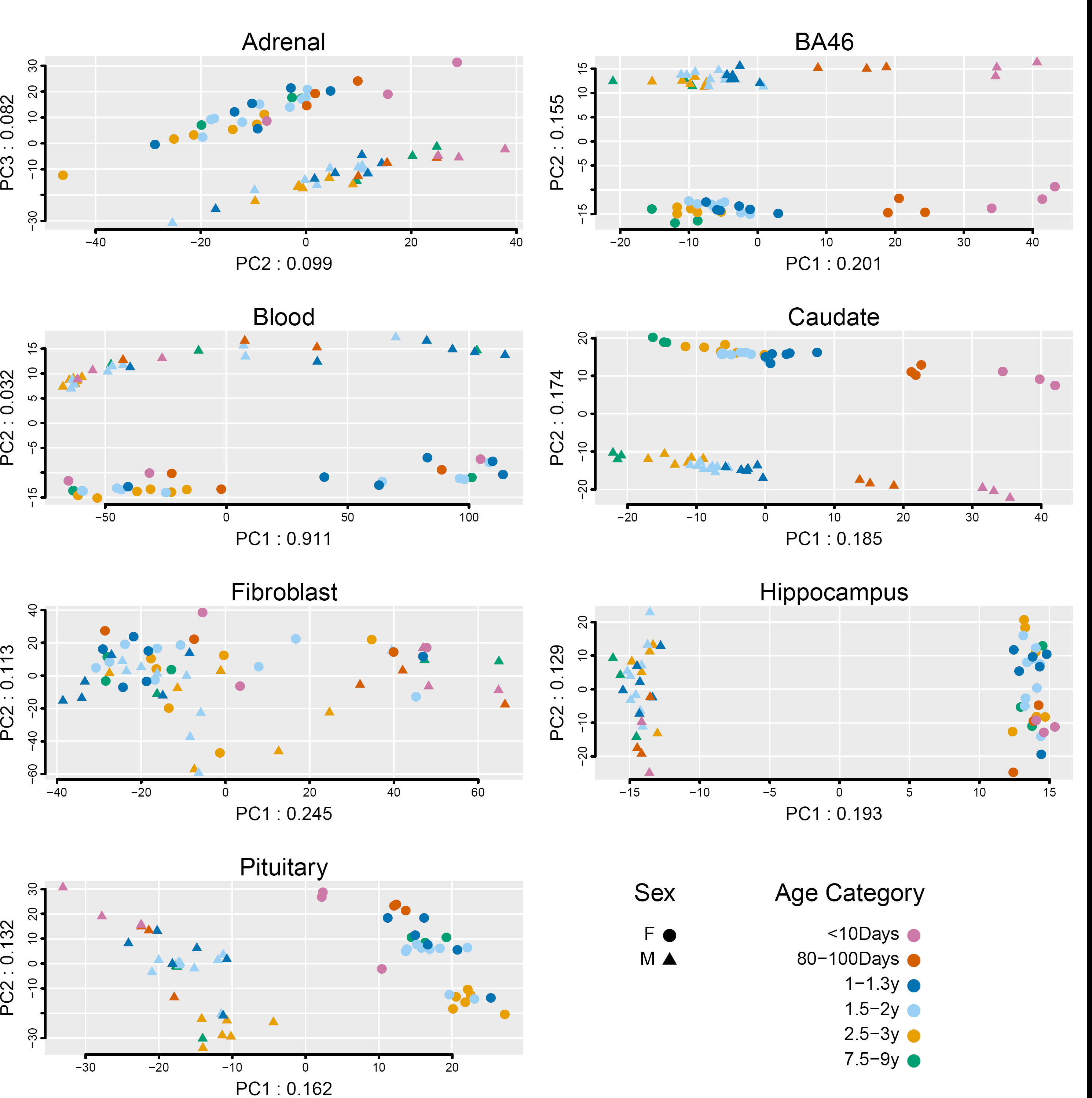
PCA of 1,000 genes with the most variable expression levels. Analysis was performed separately by tissue; sample size was 60 animals for adrenal, blood, fibroblasts, and pituitary and was 59 for BA46, caudate, and hippocampus. Numbers in the labels for x and y axes indicate the proportion of total variance accounted for by that PC.

Among the seven vervet tissues, the patterns in BA46 and caudate display the clearest association with development; PC1 (20.1% of BA46 variability and 18.5% of caudate variability) distinguishes the vervets in a nearly linear manner, with increasing age. All tissues except fibroblast show a sharp demarcation in expression pattern between males and females; this differentiation is observed on PC1 for hippocampus and pituitary (19.3% and 16.2% of variability, respectively), on PC2 for BA46, caudate and blood (15.5%, 17.4%, and 3.2% of variability, respectively), and on PC3 for adrenal (8.2% of variability).

As an initial, descriptive exploration of the biology underlying these tissue-related expression patterns, we identified, in the brain and endocrine tissue, the genes in the top and bottom 10% of the distribution of PC loadings on PCs 1, 2, and 3 (200 genes total per tissue, per PC). We evaluated the known functions of these genes, which contribute most to the variance explained by the PCs in relation to sex (BA46, caudate, hippocampus, pituitary, and adrenal, see Supplementary Table 7, Supplementary Material) or age (BA46 and caudate, Supplementary Table 8).

Age-related expression patterns in BA46 and caudate highlight numerous genes that are essential for nervous system development or that are implicated in human diseases. For example, three thrombospondin genes controlling synaptogenesis show a clear developmental pattern in BA46; *THBS1* and *THBS2* are upregulated in neonates, while *THBS4,* a gene upregulated during human brain evolution^18^, shows increasing expression across development (Fig. 2). Supplementary Fig. 2 illustrates striking age-related expression patterns in BA46 and caudate observed for other notable genes (see Supplementary Material). Supplementary Fig. 3 displays developmental expression profiles for the orthologs of these genes in human and rhesus macaque brain tissues that are most equivalent to vervet BA46 and caudate (Online Methods); the overall patterns are roughly similar to, but less pronounced than those we observed in vervet. Given the PCA results showing an age-related component to gene expression variation that differs by tissues, we conducted a differential expression analysis, using age as both a continuous and a categorical predictor in two different linear models. Nearly 8,000 genes across all seven tissues show significant differential expression by age for either analysis, mostly with very small effects (Supplementary Table 9)

**Fig. 2.**
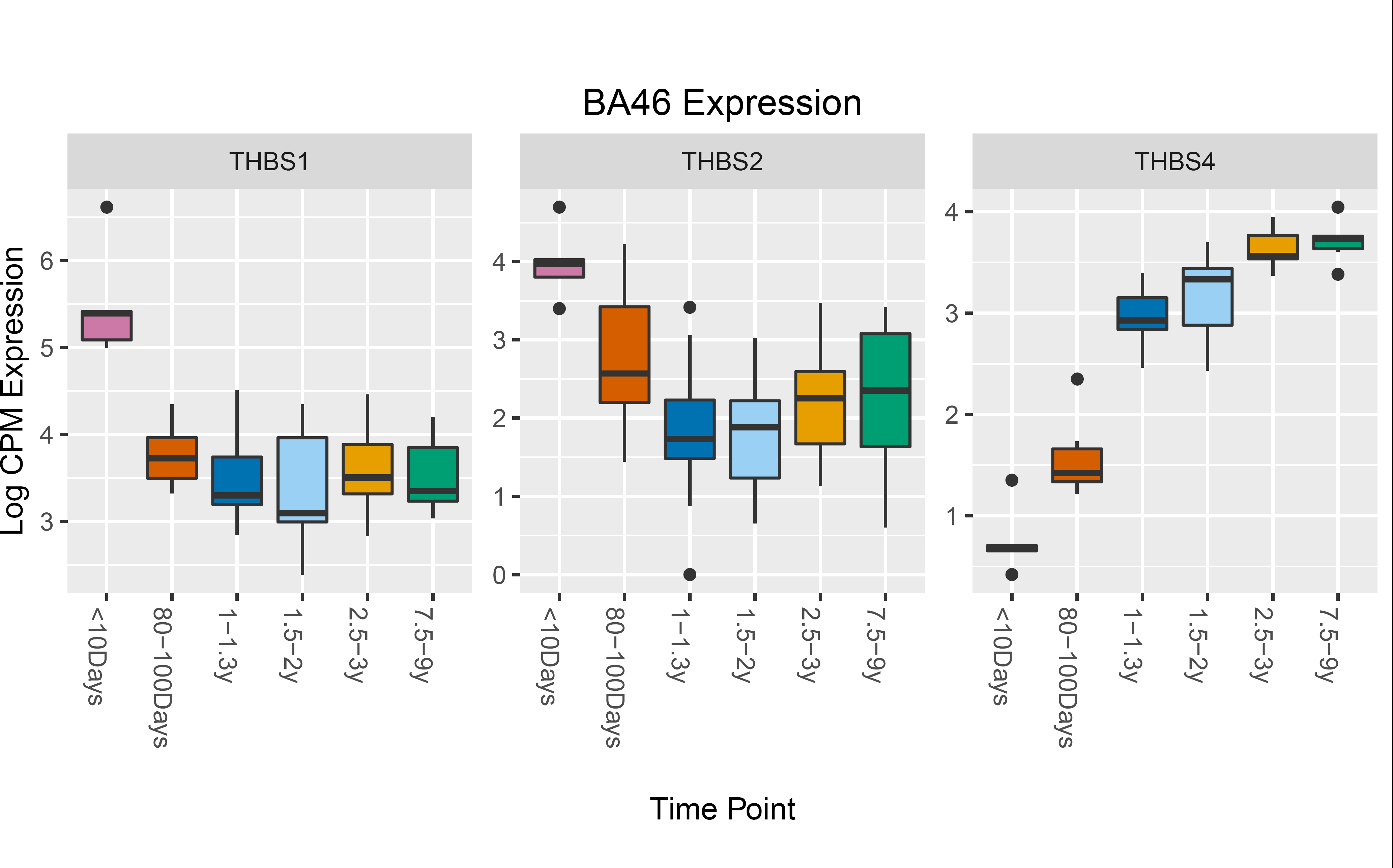
Boxplot of log counts per million (CPM) expression in samples of BA46 from 58 animals vs. timepoint, for three genes with a strong relationship between expression pattern and age. The inter-quartile range defines the height of the box, and whiskers extend to 1.5x the inter-quartile range. Outliers are indicated as individual points. In each box, the median is represented by the horizontal black bar.

**Fig. 3.**
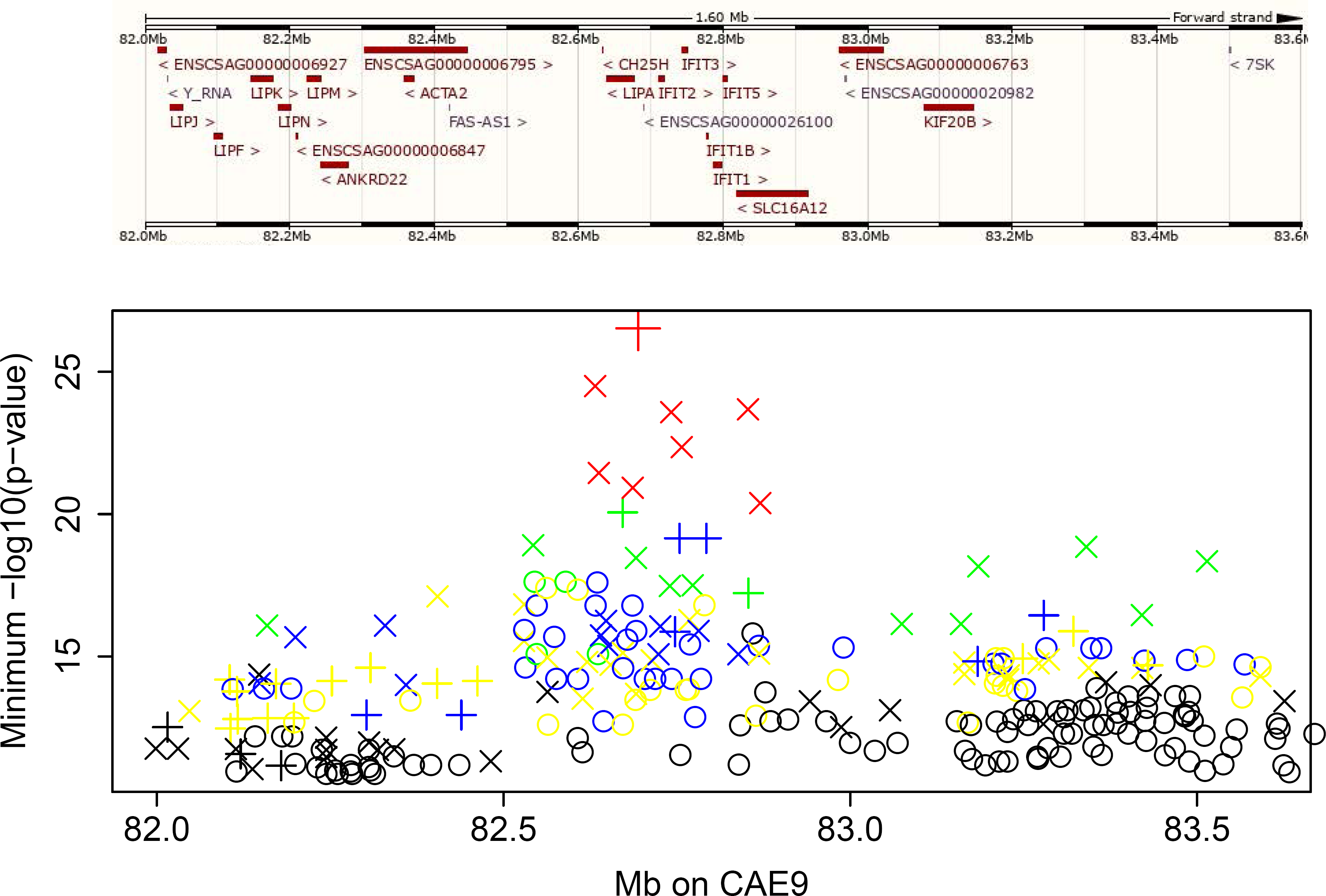
Master regulatory locus on vervet chromosome CAE 9. Upper panel: Ensembl view of the CAE 9 region. Lower panel: The minimum −log10(p-value) for each SNP in association analyses vs. expression in 347 animals of microarray probes on different chromosomes. The symbols are color-coded to represent the number of probes significantly associated to each SNP: 1-2 probes (black), 3-4 probes (yellow), 5-6 probes (blue), 7-10 probes (green), 11-14 probes (red). Symbols indicate the p-value from analysis of expression in Dataset 2 (RNA-Seq). Cross: p<2.35e-05; X: p<0.001; circle: p>0.001. The large red X at the top of the plot is CAE9_82694171.

We considered that cell-type heterogeneity could influence the interpretation of our expression and eQTL results, particularly for blood and the three brain tissues. To evaluate such heterogeneity we conducted a transcriptional deconvolution analysis of these tissues, using published data^19,20^ (Supplementary Fig. 4-7). We estimated the diversity of cell types per sample in each tissue by calculating entropy and observed that blood has substantially higher diversity of cell types than do the three brain tissues (Supplementary Fig. 8).

We also examined the relationship between the proportion of specific cell types and developmental stage. For BA46 and hippocampus, the proportion of Oligodendrocyte Precursor cells decreases as age increases, which is consistent with data from a prior study in human^21^, while the proportion of this cell type in caudate increases with increasing age. Similarly, the proportion of neurons increases as age increases in BA46 and hippocampus, and decreases with increasing age in caudate. (Supplementary Fig. 4-6). We found no correlation between estimated cell proportions and major PC axes in any tissue.

We evaluated the potential impact of technical variables on transcriptomic profiles and PC patterns (Supplementary Material). RNA-Seq sample batch demonstrated an association with expression profiles in pituitary and adrenal (PC2) and caudate and pituitary (PC3); we therefore included batch as a covariate in eQTL analysis.

### Identification of eQTLs

Whole genome sequencing (WGS) of 721 VRC monkeys has previously provided the first NHP genome-wide, high-resolution genetic variant set^11^: 497,163 WGS-based SNPs that tag common variation genome-wide. Using these SNPs we conducted separate GWAS of Datasets 1 and 2 to identify local (probes/genes < 1 Mb from an associated SNP) and distant (all other probe/gene-SNP associations) eQTLs in each dataset. Covariates in all eQTL analyses included age, sex, and batch.

We used SOLAR^22^ to estimate heritability of probe expression in Dataset 1, identifying significant heritability for 3,417 probes (out of the 6,018 filtered probes that we evaluated, corresponding to 5,586 unique genes) at a false discovery rate (FDR) threshold < 0.01 (Supplementary Data 1, 2). In a GWAS of each heritable probe, we identified 461 local and 215 distant probes to have one or more eQTLs (significant at Bonferroni-corrected thresholds of 4.8 × 10^-8^ for local and 1.5 × 10^-11^ for distant eQTLs, Table 1, Supplementary Data 3). Approximately 35% of probes with a significant eQTL (173/498) displayed at least one local *and* one distant significant association.

**Table 1.** Gene expression data sets. The number of probes/genes with at least one significant local and distant eQTL (at Bonferroni corrected thresholds) are presented. We have 80% power to detect distant eQTLs accounting for 15% of the variability in expression in Dataset 1 and 66% of the variability in Dataset 2

| Tissue | Probes/genes analyzed<sup>a</sup> | Local eQTL<sup>b</sup> | Distant eQTL<sup>c</sup> | %Distant eQTL on same chr |
| --- | --- | --- | --- | --- |
| **Dataset 1:**<br>**Microarray** |  |  |  |  |
| Blood | 3,417 | 461 | 215 | 80.8% |
| **Dataset 2:**<br>**RNA-seq** |  |  |  |  |
| Adrenal | 25,187 | 555 | 80 | 54.5% |
| BA46 | 27,530 | 307 | 30 | 81.8% |
| Blood | 33,776 | 60 | 4 | 100% |
| Caudate | 28,249 | 441 | 47 | 69.0% |
| Fibroblast | 22,328 | 239 | 43 | 33.2% |
| Hippocampus | 26,957 | 361 | 45 | 70.6% |
| Pituitary Gland | 27,236 | 596 | 80 | 77.5% |

<sup>a</sup>microarray dataset (Dataset 1) with an initial set of 22,184 probes on Illumina HumanRef-8 v2 (6,018 probes passed filters described in Supplementary Table 1; 3,417 were heritable); RNA-seq (Dataset 2) with an initial set of 33,994 genes annotated in vervet
<sup>b</sup>Local eQTL are eQTL that are within 1 Mb of the gene. Bonferroni threshold for Dataset 1: $4.8 \times 10^{-8}$ ; Bonferroni threshold for Dataset 2: $6.5 \times 10^{-10}$
<sup>c</sup>Distant eQTL are more than 1 Mb away from the gene, and may be on the same or a different chromosome. Bonferroni threshold for Dataset 1: $1.5 \times 10^{-11}$ ; Bonferroni threshold for Dataset 2: $5.3 \times 10^{-13}$

In Dataset 2 we observed, for each of the five solid tissues, between 361-596 genes with local eQTLs and 30-80 genes with distant eQTLs, and for blood and fibroblasts, 60 and 239 genes with local eQTLs and 4 and 43 genes with distant eQTLs, respectively, all at Bonferroni corrected significance thresholds (6.5 × 10^-10^ [local] and 5.3 × 10^-13^ [distant]) (Table 1, Supplementary Data 4). The smaller number of eQTLs observed in blood likely reflects heteroegentiy in the proportions of different cell types in this tissue as identified in deconvolution analyses (Supplementary Fig. 1, 8); we have no obvious explanation for the relative paucity of eQTLs in fibroblasts, aside from the observation that fewer genes were analyzed in fibroblasts than in tissues with cellular heterogeneity. At Bonferroni significance levels, we had 80% power to detect a significant local eQTL accounting for 11% of variability in expression in Dataset 1, and accounting for 55% of variability in expression in Dataset 2. For about 70% of Bonferroni-significant eQTLs (local and distant and in all tissues), the SNPs demonstrating association had minor allele frequency > 30% (Supplementary Table 10).

We considered the possibility that genotypic variation within the vervet pedigree could confound the effects of age in generating the strong loadings on PCs associated with age in BA46 and caudate. Among the 200 genes with such strong loadings, 26 of 200 genes in BA46 showed evidence of an eQTL, and for only one gene (*LOC103219658*) could genotype partially account for the association with age. Similarly, 37 genes showed evidence of eQTLs in caudate, even when using the more liberal FDR controlling procedure. For these 37 genes, we modeled expression as a function of both age and genotype, using the most significant eQTLs, and found that genotype could not account for the association with age (data not shown).

We evaluated the enrichment/depletion of cis-eQTLs in genes with age effects, using genes without age effects as reference (Supplementary Table 9). We observe that the genes with age related pattern are actually depleted for eQTLs (Supplementary Table 11), in accord with prior studies predicting that purifying selection results in such depletion in genes that play important roles at particular developmental timepoints^23^.

### Comparison to Human eQTLs

While the eQTLs summarized in Table 1 exceeded Bonferroni thresholds, we also applied FDR-controlling procedures, to expand the list of local eQTLs for more exploratory investigations, and to make our results comparable to those of GTEx (Table 2). We controlled the FDR for eGenes at 0.05 (Online Methods), accounting for multiple testing using a hierarchical error controlling procedure developed for multi-tissue eQTL analysis^24^. We applied this same procedure to GTEx eQTLs to facilitate the comparison between the datasets.

**Table 2.**
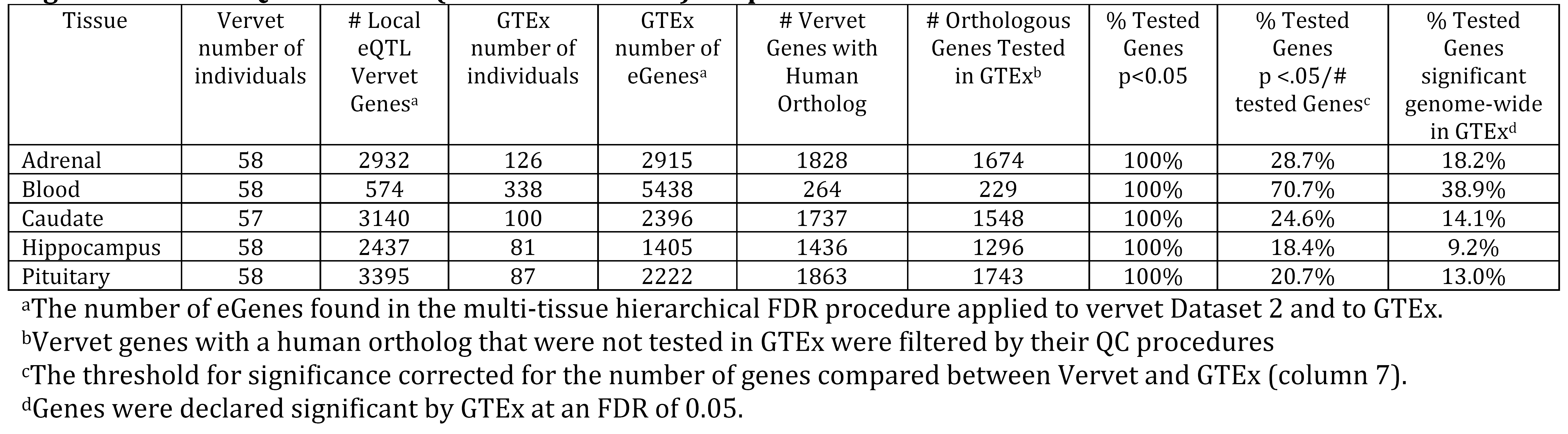
Comparison of specific genes with local eQTL in Vervet Dataset 2 to GTEx. The number of genes with at least one significant local eQTL in Vervet (at FDR thresholds) are presented.

| Tissue | Vervet number of individuals | # Local eQTL Vervet Genes<sup>a</sup> | GTEx number of individuals | GTEx number of eGenes<sup>a</sup> | # Vervet Genes with Human Ortholog | # Orthologous Genes Tested in GTEx<sup>b</sup> | % Tested Genes p<0.05 | % Tested Genes p <.05/# tested Genes<sup>c</sup> | % Tested Genes significant genome-wide in GTEx<sup>d</sup> |
| --- | --- | --- | --- | --- | --- | --- | --- | --- | --- |
| Adrenal | 58 | 2932 | 126 | 2915 | 1828 | 1674 | 100% | 28.7% | 18.2% |
| Blood | 58 | 574 | 338 | 5438 | 264 | 229 | 100% | 70.7% | 38.9% |
| Caudate | 57 | 3140 | 100 | 2396 | 1737 | 1548 | 100% | 24.6% | 14.1% |
| Hippocampus | 58 | 2437 | 81 | 1405 | 1436 | 1296 | 100% | 18.4% | 9.2% |
| Pituitary | 58 | 3395 | 87 | 2222 | 1863 | 1743 | 100% | 20.7% | 13.0% |

<sup>a</sup>The number of eGenes found in the multi-tissue hierarchical FDR procedure applied to vervet Dataset 2 and to GTEx.
<sup>b</sup>Vervet genes with a human ortholog that were not tested in GTEx were filtered by their QC procedures
<sup>c</sup>The threshold for significance corrected for the number of genes compared between Vervet and GTEx (column 7).
<sup>d</sup>Genes were declared significant by GTEx at an FDR of 0.05.

In comparison with GTEx V6, despite having a smaller sample size we identify more local eQTLs (at FDR thresholds applied to both datasets, see Online Methods) for the five solid tissues that were evaluated in both resources (Table 2). We attribute the larger number of local eQTLs identified in the vervet sample, relative to GTEx, to the more homogenous environment of colonied NHPs compared to humans, and to the more uniform process of collecting tissues in this study. We also evaluated the degree to which specific vervet and GTEx eQTLs overlap. All genes with a genome-wide significant vervet eQTL (at FDR <0.05) also display a human eQTL in the same tissue (at p< 0.05), given that the gene has a known human ortholog and was tested in GTEx. Using instead, GTEx’s defined significance threshold for orthologous genes (FDR < 0.05), an average of 19% of vervet eQTLs display such a human eQTL (Table 2). Restricting the comparison to Bonferroni-significant local eQTLs, an average of 23% of vervet eQTLs also have such an eQTL in the same tissue in GTEx (Supplementary Table 12).

We also compared our local eQTL results for brain tissues to the Open Access version of human eQTLs from DFPLC, available from CommonMind Consortium (CMC)^25^. More than 87% of vervet brain local eQTL genes with human orthologs in the CMC dataset have a local eQTL at FDR<0.05 in that dataset (Supplementary Material and Supplementary Table 13).

### eGene Sharing Among Tissues

We assessed sharing of locally regulated eGenes (genes with a significant local eQTL, see Online Methods) across tissues (Supplementary Fig. 9). We differentiated between tissue-specific and shared eGenes. We observed that the tissue-specific eGenes in all tested tissues except blood are more common than eGenes shared among tissues. The largest number of shared local eGenes was observed between adrenal and pituitary (300), organs inter-regulated in the same neuroendocrine pathway, and then among the three brain regions (239); 229 eGenes are shared across all tissues but blood and 82 eGenes are shared across all seven tissues.

### Genomic Distribution of eQTLs

Regulatory variants occur most frequently in functional genomic regions^26^, and we find that vervet gene regions encompassing exons, introns and adjacent flanks show a clear enrichment for local eQTLs (Supplementary Fig. 10, Supplementary Table 14). Conversely, intergenic regions show a significant deficit of local eQTLs (Supplementary Fig. 10, Supplementary Table 14). As in other primates^27^, vervet eQTLs are enriched around gene boundaries (transcription start site [TSS] and transcription end site [TES]) (Supplementary Fig. 11).

We used previously published chromatin immunoprecipitation with DNA sequencing (ChIP-Seq) data^28,33^ to evaluate eQTL distribution in H3K4me3 enriched regions (promoters) and H3K27ac enriched regions (which include acetylated promoters and enhancers). As H3K4me3 marks are typically conserved across tissues we analyzed them using vervet liver data. As enhancer marks are more tissue specific^29-31^ we analyzed H3K27ac marks in both vervet liver and available brain data (caudate and prefrontal cortex) from rhesus macaque^28^. The promoter regions show stronger enrichment for vervet local eQTLs than either genic or H3K27ac-enriched regions (Supplementary Fig. 10, Supplementary Table 14).

### Validation of Distant eQTLs: a Master Regulatory Locus on Vervet Chromosome 9

Our Dataset 1 is well-powered for discovery of distant eQTLs. Among the 215 genes for which we observed association at genome-wide significance thresholds to one or more distant eQTLs, a locus on CAE9 in which 76 SNPs across a ~500 Kb region displayed genome-wide significant local eQTL signals, stood out for showing association to multiple unlinked genes. For each of these 76 SNPs we identified genome-wide significant distant eQTLs at between five and 14 genes, on different vervet chromosomes, for a total of 2,127 distant SNP-gene associations (Fig. 3, Supplementary Table 15).

Because we obtained Dataset 2 using a different platform from Dataset 1, and from a mostly non-overlapping sample (only 6 vervets were in both datasets), we evaluated it for replication of the CAE 9 distant eQTLs, recognizing the limited power of this much smaller dataset. Considering the percent of variance accounted for by the distant eQTLs in Dataset 1 (Supplementary Table 15), we have 82% power to identify eQTLs in Dataset 2, with 58 animals, when the SNP accounts for 35% or more of expression variance, using a significance threshold (p<2.35 × 10^-5^) that accounts for multiple testing of the 76 SNPs to multiple genes (2,127 tests). Two genes, *ST7* (31 SNPs) and *YPEL4* (22 SNPs) replicate association at this threshold, with estimated regression coefficients for these 53 SNP-gene associations being similar in magnitude and direction in the two datasets (Supplementary Table 16). We confirmed eight distant associations (*RANBP10, LCMT1, ST7, TMEM57, YPEL4, NARF, STXBP1, DEDD2*) across the two datasets, with at least one SNP demonstrating association at a marginal p<0.05 (Supplementary Table 15).

These results suggest that the CAE 9 eQTL represents a master regulatory locus (MRL). This genomic segment contains a cluster of acid lipase genes and interferon-inducible genes, including *IFIT1B* (Interferon-Induced Protein With Tetratricopeptide Repeats 1B), a gene recently implicated in viral resistance in vervets, but not humans^32^. The same SNPs contributing to the MRL are also local eQTLs for *IFIT1B,* at genome-wide significant levels, however GTEx reports no significant local eQTLs for *IFIT1B* in human blood.

Expression of *IFIT1B* correlates strongly with expression of the distant genes regulated by this eQTL (Supplementary Material, Supplementary Table 17). We conducted mediation analyses in Dataset 1 for a SNP (CAE9_82694171) that, at Bonferroni corrected significance thresholds, is both a distant eQTL for all 14 genes and a local eQTL for *IFIT1B* (Supplementary Table 18). This SNP accounts for 19-37% of the variance in expression level of the 14 genes not located on CAE 9. When we conditioned these analyses on expression of *IFIT1B,* the magnitude of these distant associations diminished substantially, the variance accounted for by this SNP dropping to 10% or less for all 14 genes. These results indicate that *IFIT1B,* under direct control of a local eQTL on CAE 9, likely influences expression of 14 other genes spread across the genome. As suggested by studies in human populations, such phenomenon of mediation by local eQTLs of distant eQTLs provides a further validation of the latter loci^33^.

### Identification of Hippocampus-Specific eQTLs in a Region Linked to Hippocampal Volume

In an initial investigation of the impact of vervet tissue-specific eQTLs on higher order traits we focused on MRI-based hippocampal volume, a highly heritable trait in the VRC (h^2^ =0.95)^34^, for which the strongest QTL signal genome wide (peak LOD score 3.42) lies in an ~8.3 Mb segment of CAE 18. Power simulations in SOLAR indicate that, in the VRC pedigree, quantitative trait data for 347 vervets (the number with hippocampal volume data) provide 80% power to detect a locus with LOD=2 when locus-specific heritability is > 45%. In the center of the broad region around this linkage peak, two hippocampus-specific local eQTLs were Bonferroni-significant at a genome-wide threshold (Fig. 4).

**Fig. 4.**
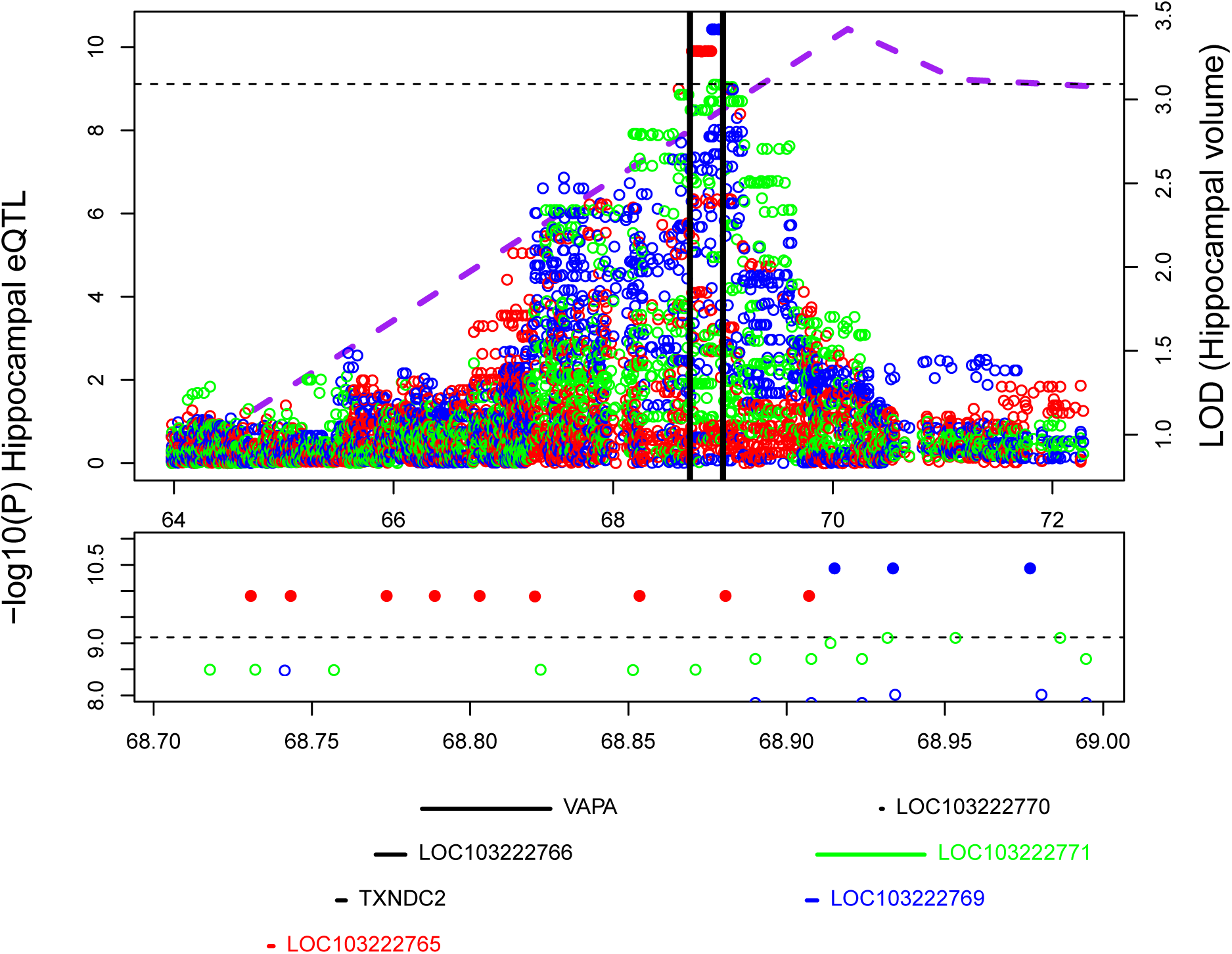
Hippocampal volume QTL and local hippocampal volume eQTLs in RNA-Seq analysis.Top panel: purple dotted line is the multipoint LOD score for hippocampal volume (measured in 347 animals). Circles represent evidence for association of SNPs to hippocampal expression in 58 animals of three genes: *LOC103222765* (red), *LOC103222769* (blue) and *LOC103222771* (green). Solid circles indicate genome-wide significant associations. The region between the black vertical lines is blown up in the middle and bottom panels. The horizontal dotted line represents the genome-wide significant threshold for local eQTLs. Middle panel: SNPs with −log10(p-value)>8 for association to expression in hippocampus, color codes are as in the top panel. Bottom panel: Genes sited between 68.7 and 69 Mb (the eQTL region). Color codes are as in the top panel.

The genome-wide significant eQTLs SNPs reside in, and regulate expression of, two lncRNAs located 168 Kb apart: *LOC103222765* (nine associated local eQTL SNPs) and *LOC103222769* (three associated local eQTL SNPs). An additional lncRNA gene, *LOC103222771,* situated two bp from *LOC103222769,* shows hippocampal specific association to six SNPs at a significance level (p < 10^-9^) just above the genome-wide Bonferroni-corrected threshold. While all three genes display hippocampus-specific eQTLs, the genes themselves are expressed across all seven tissues that we analyzed, and show no significant sex or age specific differences in expression patterns (data not shown). The incomplete database annotation of lncRNAs^35^ limits comparative analyses of such genes among primates; a BLAST search found a homolog for *LOC103222765* in the white-tufted-ear marmoset and one for *LOC103222771,* in the crab-eating macaque. While *LOC103222765* overlaps a coding gene (*RAB31*), *LOC103222769* and *LOC103222771* do not overlap exons of any coding genes and therefore are more specifically classified as long intergenic non-coding RNA (lncRNA) genes^36^.

Given the physical proximity of these lncRNAs, we used multivariate conditional analyses to evaluate whether the regulation of these genes depends on a single or multiple independent eQTLs. For each lncRNA we designated a “lead SNP” (the SNP most significantly associated to its expression, Supplementary Table 19). For both *LOC103222769* and *LOC103222771,* modeling expression as a function of both lead SNPs results in diminished significance levels for both SNPs (Supplementary Table 19), suggesting that one eQTL regulates both genes. Modeling *LOC103222765* expression as a function of its lead SNP and the lead SNP of the other two genes, the lead SNP for *LOC103222765* remains significant, while the other two lead SNPs are non-significant, confirming the “distinctness” of this signal (Supplementary Table 19). This analysis suggests two eQTLs in this region; one associated with *LOC103222765,* and the second associated with *LOC103222769* and *LOC103222771.*

We observed a positive correlation between hippocampal expression of *LOC103222765, LOC103222769* and *LOC103222771,* and hippocampal volume as assessed by MRI, in six vervets for which both MRI and RNA-Seq data were available. To extend this observation, we assessed, using an independent platform, quantitative real-time PCR (qRT-PCR), *LOC103222765, LOC103222769* and *LOC103222771* hippocampal expression in these six vervets and 10 additional vervets for which both hippocampal RNA and MRI data were available. In this expanded sample set, we identified significant positive correlations (Fig. 5) between *LOC103222765, LOC103222769* and *LOC103222771* expression and hippocampal volume. While the above data suggest that genetic variation in this region regulates these lncRNAs and also has a strong impact on the MRI phenotype, colocalization analysis^37^ does not support the hypothesis that a single variant accounts for both the genome-wide linkage (MRI) and GWAS (eQTL) findings (8.2% posterior probability).

**Fig. 5.**
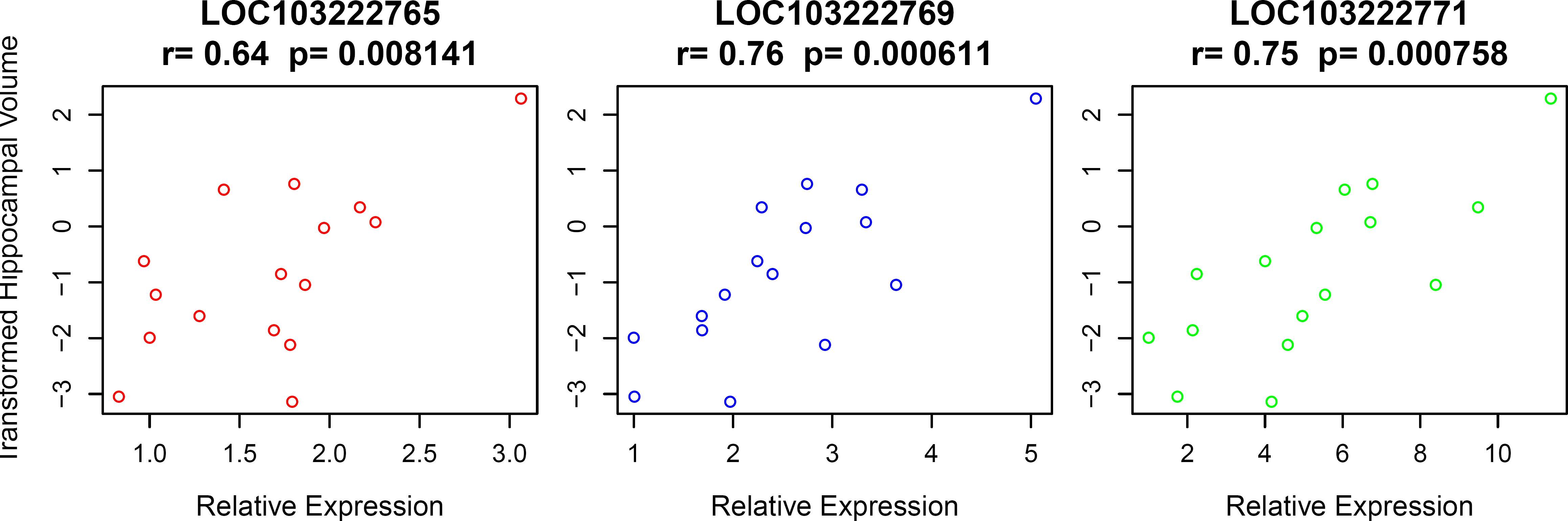
Correlation in 16 animals of hippocampal volume (MRI) with hippocampal expression of *LOC103222765* (left), *LOC103222769* (middle) and *LOC103222771* (right). The expression data are from qRT-PCR. Quantification was performed using the relative standard curve method, with the geometric mean of the reference gene *HPRT1* used as an endogenous control for normalization of the interpolated lncRNA quantities. Hippocampal volume measurements are residuals from a regression on covariates of age and sex. “r” is the Pearson correlation coefficient, and the p-value tests the null hypothesis that r=0.

## Discussion

The data presented here provide the first NHP resource for investigating the genetic contribution to inter-individual variation in gene expression across multiple tissues and development. This resource, in a species closely related to humans, complements GTEx, which has become an essential tool for pinpointing genes, and even variants, underlying human GWAS findings^38,39^.

Several features differentiate this vervet resource from GTEx, reflecting aspects of the study design that are infeasible in human research. Notably, the age-based sampling design enabled us to delineate tissue-specific expression profiles in relation to developmental trajectories. Delineating these trajectories provides insights into biological processes that may be associated with the expression profiles of particular genes. For example, several genes that contribute to synapse formation and postnatal myelination of the central nervous system^40-43^ contribute to the near linear age-related pattern observed in BA46 and caudate and, and suggest that the observed expression pattern reflects this process. Conversely, the lack of such a developmentally specific pattern in the hippocampus may relate to the generation of functional neurons in this tissue that occurs throughout the lifespan, underpinning its functions in learning and memory^44,45^.

Three factors increased the signal-to-noise ratio of vervet eQTL analyses, relative to human studies: (i) the homogeneity of the vervet sample with respect to environmental exposures; (ii) the greater control over necropsy conditions; and (iii) the restricted genetic background of the recently bottlenecked Caribbean vervet population. These factors enabled us to identify 385 genes with one or more genome wide significant distant eQTLs, including the MRL at *IFIT1B.*

The function of *IFIT1B,* one of a cluster of five IFIT genes, is poorly understood. It is a paralog of *IFIT1,* which is involved in innate antiviral immunity in mammals, broadly^46^, and in regulation of gut microbiota in mouse^47^. In some mammalian species *IFIT1B* contributes to discrimination between “self versus non-self” transcripts based on the lack of 2’ O-methylation on mRNA 5’ caps in viruses, a so-called cap0 structure^32^. Vervet *IFIT1B* recognizes and inhibits replication of viruses with cap0-mRNAs, while human *IFIT1B* lacks this function^32^. This functional divergence of *IFIT1B* antiviral activity may reflect the divergence of the human lineage from that of other primates, in exposures and adaptations to particular pathogens, including the arboviruses which are responsible for diseases such as encephalitis, dengue, and yellow fever.

Our results suggest that investigation of genes regulated by *IFIT1B* in vervet might reveal mechanisms for its role in defense against viral pathogens. While these genes do not act together in any annotated pathway, recent evidence points to immune functions for the products of several of them. For example, *RANBP10,* a transcriptional coactivator, promotes viral gene expression and replication in HSV-1 infected cells^48^. *SUGT1,* a cell cycle regulator, is the homolog of *SGT1,* which plays an essential role in innate immunity in plants as well as mammals^49,50^, while *TMEM57* shows genome-wide significant association in human to blood markers of inflammation^51^.

Just as GTEx data are helping refine signals from human GWAS of complex traits^5^, we used vervet hippocampal eQTLs to identify a set of lncRNAs as candidate genes for a higher order phenotype, hippocampal volume. The genetic and environmental homogeneity of the relatively small vervet study sample likely facilitated these findings, and supports the extension of multi-tissue vervet eQTL studies as a strategy for identifying loci with a large impact on higher-order phenotypes, generally. As the tissues examined to date are only a fraction of those available from the same set of vervets, it will be possible to extend the investigations reported here to an additional 60 brain regions and 20 peripheral tissues.

While expanding expression resources in other NHP species will create additional opportunities to identify eQTLs that are informative for various biomedical investigations^9,52^, the Caribbean vervet is unique among NHPs in having abundant natural populations available for such investigations, with an essentially identical genetic background to the samples studied here^10,13^. For example, the lead SNPs for the eQTLs contributing to hippocampal volume in the VRC each occur at a relatively high frequency in these island populations (Supplementary Material). We therefore anticipate that most findings presented here can be followed up through well-powered association studies.

## Online Methods

### Study Sample

The vervet monkeys used in this study are part of the Vervet Research Colony (VRC), established by UCLA during the 1970’s and 1980’s from 57 founder animals captured from wild populations in St. Kitts and Nevis^10^. In 2008 the VRC was moved to Wake Forest School of Medicine; the MRI phenotypes included in this study were collected when the colony was in California (see Supplementary Material for more details). All of the animals in this study were captive-born, mother-reared and socially-housed in large, indoor-outdoor enclosures, in matrilineal groups that approximated the social structure of wild vervet populations. They had a uniform exposure to light and darkness and were fed a standardized diet.

### Gene Expression Phenotypes

Two data sets of gene expression measurements were collected. Dataset 1 consisted of microarray (Illumina HumanRef-8 v2) assays of RNA obtained from whole blood in 347 vervets, while Dataset 2 consisted of RNA-Seq data from 60 animals, with seven tissues assayed in each animal. Six vervets were in both Datasets; no randomization was applied in allocating animals to Datasets and investigators were not blinded to the allocation of animals to Datasets.

#### Dataset 1: Microarrays From Whole Blood

The microarray data set has been described in Jasinska et al.^13^ and is available at NCBI at the BioProject PRJNA115831. Details on RNA extraction, cDNA synthesis, and initial data processing are presented in Supplementary Material. To obtain a set of probes usable in vervet from the Illumina HumanRef-8 v2 microarray (originally developed for assaying gene expression in humans), we used the vervet reference sequence to select probes that contain no vervet indels and demonstrate ≤ five mismatches, with a maximum of one mismatch in the 16 nt central portion of the probe. To prevent bias in the measurement of expression due to SNP interference with hybridization, we excluded probes targeting sequences with common SNPs identified in the VRC pedigree. A total of 11,001 probes passed these filters (Supplementary Table 1). Illumina provides a “detection p-value” for each subject and probe; p<0.05 indicates significant detection of a given probe in a specific individual. We retained for analysis 6,018 probes that were detected with detection p-values of p<0.05 in at least 5% of vervets, and tested for association 3,417 probes that were significantly heritable. Expression data were inverse-normal transformed prior to analysis.

#### Dataset 2: RNA-Seq Data from Seven Tissues

Tissues harvested during experimental necropsies were obtained from 60 vervets representing 10 developmental stages, ranging from neonates (7 days), through infants (90 days and one year), young juveniles (1.25, 1.5, 1.75, 2 years old), subadults (2.5, 3 years old) to adults (4+ years old), with six vervets (3 male and 3 female) from each developmental time point. The IACUC protocol number covering the necropsies was A09-512. This necropsy protocol was approved by the IACUC at Wake Forest School of Medicine. Two vervets (a 1.75 year old female and a 7 day old male) for which we did not have WGS data were excluded from the eQTL study. Altogether, in the eQTL study we included 11 vervets below one year old, 23 vervets between one to two years old, and 24 vervets between two and four years old, 29 males and 29 females. Details regarding tissue collection and RNA collection procedures are in Supplementary Material.

We conducted RNA-Seq for all vervets in seven tissues: three brain tissues (BA46, caudate and hippocampus), two neuroendocrine tissues (adrenal and pituitary) and two peripheral tissues serving as a source of biomarkers (blood and fibroblasts). From purified RNA, we created two types of cDNA libraries, poly-A RNA (fibroblasts, adrenal and pituitary) and total RNA (blood, caudate, hippocampus, BA46) cDNA libraries (Supplementary Table 20, Supplementary Material). For one vervet the RNA-Seq data indicated that the caudate and BA46 samples had been mixed-up, and for this vervet we therefore did not include the data from these two tissues in any analyses. Details on library preparation are in Supplementary Material. The RNA-Seq read data were made available through NCBI as BioProject PRJNA219198.

RNA-Seq reads were aligned to the vervet genomic assembly Chlorocebus_sabeus 1.1 http://www.ncbi.nlm.nih.gov/assembly/GCF000409795.2 by the ultrafast STAR aligner^53^ using our standardized pipeline. STAR was run using default parameters, which allow a maximum of ten mismatches. Gene expression was measured as total read counts per gene. For paired end experiments, total fragments are considered. Fragment counts that aligned to known exonic regions based on the NCBI *Chlorocebus sabaeus* Annotation Release 100 were quantified using the HTSeq package^54^. The counts for all 33,994 genes were then combined, and lowly expressed genes, defined as genes with a mean in raw counts of < 1 across all samples, as well as genes detected in fewer than 10% of individuals were filtered out. The calcNormFactors function in the edgeR package^55^ was applied to normalize counts. Finally, an inverse-normal transform was applied to counts per million prior to analysis.

Deconvolution analysis was perfomed in vervet brain and blood tissue using available reference for these tissues. For the brain tissues, gene signatures were obtained from Zhang et al.^20^, for blood, cell type specific markers were taken from datasets built into the CellMix package^19^. Cell type composition for each tissue was evaluated using the CellMix R package.

#### Datasets for comparative expression analysis between species

We performed comparative analysis of gene expression between vervet brain regions and age-matched human and rhesus macaque samples. We compared overall expression profiles between these species and inspected developmental expression patterns of selected genes.

We paired age categories between vervet and two primate species with developmental gene expression data available from the Allen Brain Atlas (ABA). Gene expression in human from BrainSpan dataset was assessed using RNA-Seq, and gene expression in rhesus macaque from the NIH Blueprint Non-Human Primate (NHP) Atlas was assessed using microarray^6,52^ (Supplementary Tables 21, 22). We matched the three vervet brain tissues to the most closely corresponding available tissues in the two other species (Supplementary Table 23).

Overall mean levels of expression were compared between species using a rank correlation. For the comparison with human, two independent analyses were performed using two different datasets: GTEx data and ABA developmental data. The rhesus macaque comparison was limited to a single developmental dataset of male animals, also obtained from the ABA. Analyses involving the ABA developmental datasets were limited to the three brain regions most closely related to the brain tissues analyzed in vervets (Supplementary Table 23). For the GTEx comparison, vervet tissues were matched to the five corresponding tissues available: adrenal, blood, caudate, hippocampus and pituitary. As the ABA rhesus macaque dataset included only males, we limited comparisons to male vervets.

For each of the three dataset comparisons, vervet raw counts were first converted to RPKM values using the edgeR R package^55^. GTEx and human ABA counts obtained were already normalized to RPKM values; rhesus macaque counts had been normalized using an RMA approach^52^. Mean expression was then calculated by tissue for vervet and comparison datasets. For comparisons to ABA developmental datasests, mean expression was calculated by tissue type and time point, according to matched age groups (Supplementary Tables 21, 22). Vervet gene names were converted to their corresponding human orthologs to ensure gene names matched between vervet and comparison datasets; Genes with no human ortholog were excluded. In addition, genes not present in both vervet and the comparison species dataset were also removed. Variances were then calculated for each gene across the five or three different vervet tissues, for GTEx and ABA comparisons, respectively. The top 1,000 genes with the highest variances were then selected for rank-rank correlation testing. The base R function cor.test was used to perform correlation testing.

#### Real-time quantitative PCR (qPCR)

Real-time quantitative PCR was performed in two steps. First, reverse transcription (RT) was performed using the SuperScript® III First-Strand Synthesis System (Life Technologies) following the manufacturer’s protocol for priming with random hexamers. Custom primers and hydrolysis probes were designed for each lncRNA and three candidate reference genes (Hypoxanthine phosphoribosyltransferase 1, *HPRT1;* Glyceraldehyde 3-phosphate dehydrogenase, *GAPDH;* and Beta-2-Microglobulin, *B2M*) using the Custom TaqMan® Assays Design Tool (Applied Biosystems, Supplementary Table 24). Expression analyses were conducted on the LightCycler™ 480 platform (Roche) using the iTaq® Universal Probes Supermix (Bio-Rad). All qPCR reactions were carried out in triplicate and reactions containing water instead of cDNA were included as negative controls. cDNA samples were diluted 1:5 with water, and a five-point standard curve of four-fold dilutions was prepared for each gene using pooled cDNA as the template. Stability of each candidate reference gene was evaluated using the NormFinder software (v5) in R^56^. Quantification was performed using the relative standard curve method, with the geometric mean of the most stably expressed reference genes (*GAPDH* and *HPRT1*) used as an endogenous control for normalization of the interpolated lncRNA quantities. Finally, relative expression levels were generated by dividing the normalized lncRNA quantities by the corresponding quantity in one experimental sample which served as a calibrator. Refer to Supplementary Material for additional experimental details and complete primer and probe sequences information.

### Hippocampal Volume Phenotype

Estimates of hippocampal volume were measured in 347 vervets >2 years of age using MRI. Details of the image acquisition and processing protocol were described previously^34^ and are outlined in Supplementary Material. Prior to genetic analysis, hippocampal volume was log transformed, regressed on sex and age using SOLAR^22^, and residuals used as the final phenotype.

### Genotype Data

Genotype data were generated through whole genome sequencing of 725 members of the VRC^11^. Genotypes from 721 VRC vervets that passed all QC procedures can be directly queried via the EVA at EBI www.ebi.ac.uk/eva using the PRJEB7923 accession number. Two genotype data sets were used in the current study^11^: (1) The Association Mapping SNP Set consists of 497,163 SNPs on the 29 vervet autosomes. In this set of ~500K SNPs, there were an average of 198 SNPs per Mb of vervet sequence, and the largest gap size between adjacent SNPs was 5 Kb. (2) The Linkage Mapping SNP Set consists of 147,967 markers on the 29 vervet autosomes. In this set of ~148K SNPs, there were an average of 58.2 SNPs per Mb of vervet sequence, and the average gap size between adjacent SNPs was 17.5 Kb.

The software package Loki^57^, which implements Markov Chain Monte Carlo methods, was used to estimate the multipoint identical by decent (MIBD) allele-sharing among all vervet family members from the genotype data. As long stretches of IBD were evident among these very closely related animals, a reduced marker density was sufficient to evaluate MIBD at 1cM intervals; we used a 9,752 subset of the 148K SNP data set. The correspondence between physical and genetic positions in the vervet was facilitated by a vervet linkage map^58^, constructed using a set of 360 STR markers. Both the physical and genetic position of these markers was known, and genetic locations of SNPs were found by interpolation.

### Statistical Analysis

#### Principal Components Analysis (PCA)

In Dataset 2, the top 1,000 most variable genes were selected for each tissue, and PCA applied to log2-transformed counts per million, using the singular value decomposition and the prcomp function in R (https://www.R-project.org, version 3.2.3). Expression was mean-centered prior to analysis. We examined the genes in the top and bottom 10% of the distribution of PC loadings on PCs 1, 2, or 3 (200 genes total per tissue, per PC) where these loadings are taken from the eigen-decomposition of the expression matrix. The gene loadings represent the amount that gene contributes to the PC value for that sample on the axis in question.

#### Mapping of Gene Expression and Hippocampal Volume Phenotypes

We expected greater power for association analyses of gene expression traits compared to more complex phenotypes. Therefore we applied genome wide association analyses to these traits. For the higher-order phenotype examined (hippocampal volume) we anticipated having power only to detect loci with a much stronger effect, and therefore utilized linkage analysis for this trait.

##### Heritability and Multipoint Linkage Analysis

We estimated familial aggregation (heritability) of traits using SOLAR, which implements a variance components method to estimate the proportion of phenotypic variance due to additive genetic factors (narrow sense heritability). This model partitions total variability into polygenic and environmental components. The environmental component is unique to individuals while the polygenic component is shared between individuals as a function of their pedigree kinship. If the variance in phenotype Y due to the polygenic component is designated as 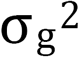 and the environmental component as 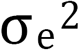, then in this model 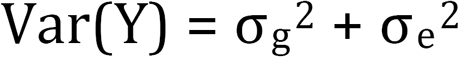, and the covariance between phenotype values of individuals *i* and *j* is 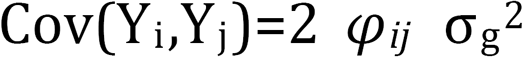 where *φ_ij_* is the kinship between individuals *i* and *j*.

Whole genome multipoint linkage analysis of hippocampal volume was also implemented in SOLAR, which uses a variance components approach to partition the genetic covariance between relatives for each trait into locus-specific heritability and residual genetic heritability. Linkage analysis was performed at 1cM intervals using the likelihood ratio statistic.

##### Association Analysis

Association between specific SNPs and gene expression phenotypes was evaluated using EMMAX^59^. EMMAX employs a linear mixed model approach, where SNP genotype is a fixed effect, and correlation of phenotype values among individuals is accounted for using an identity by state (IBS) approximation to kinship. Association analyses used the full set of 497,163 SNP markers, and for both Dataset 1 and Dataset 2 included age (where in Dataset 2 age, in days, corresponds to developmental stage), sex, and sample batch as covariates. It is common to try to account for unmeasured factors influencing global gene expression by including probabilistic estimation of expression residuals (PEER) factors as covariates^60^. We considered the controlled nature of the study environment and experimental design to preclude the need for this adjustment.

#### Colocalization of eQTL and Hippocampal Volume QTL

We evaluated the posterior probability that the hippocampal volume QTL and the hippocampus local eQTLs on CAE 18 share a single, common causal variant using COLOC^37^. The same variants were tested in both analyses, and only six vervets overlapped between the two data sets.

#### Multiple Testing Considerations in eQTL

We used a Bonferroni correction to account for multiple testing across genes, SNPs, and tissues as our primary error-controlling strategy for the identification of eQTLs. Thresholds for Dataset 2 were more stringent, as more genes were tested than in Dataset 1 (~25K vs. ~3K) and multiple tissues were analyzed in Dataset 2. Dataset 1 was analyzed association to 3,417 heritable probes. The local eQTL significance threshold (4.8 × 10^-8^) was corrected for the testing of SNPs within 1 Mb of 3,417 probes, and the distant eQTL significance threshold (1.5 × 10^-11^) accounted for genome-wide testing of 3,417 probes. Dataset 2 significance thresholds were constructed in a similar fashion, but also accounted for testing of 191,263 gene-tissue combinations (the number of genes tested per tissue is in Table 1). The RNA-Seq local eQTL threshold was 6.5 × 10^-10^, and the distant eQTL threshold was 5.3 × 10^-13^.

To identify multi-tissue eGenes, the tissues in which they are active, and the associated SNPs in each of these tissues, we used TreeBH, a hierarchical approach testing proposed in Bogomolov et al.^24^ which extends the error-controlling procedure characterized in Peterson et al.^61^ to the multi-tissue eQTL setting. To apply this method, the hypotheses are grouped into a tree with three levels: genes in level 1, tissues in level 2, and SNPs in level 3. Testing proceeds sequentially starting from the top of the tree in a manner that accounts for each previous selection step. This method allows control of the FDR of local eGenes (defined as those genes whose expression is regulated in at least one tissue by some genetic variants located within 1 Mb of the gene) and of the expected average false discovery proportion of the tissues in which we claim this regulation is present across the discovered eGenes. P-values are defined by building up from the bottom of the tree. Specifically, to obtain a p-value for the null hypothesis of no local regulation for a given gene in a given tissue (corresponding to a hypothesis in level 2 of the tree), we applied Simes’ combination rule^62^ to the p-values obtained via EMMAX for the hypotheses of no association between the expression of the gene in the tissue and each of the SNPs in the local neighborhood (corresponding to the hypotheses in level 3 of the tree). To obtain a p-value for the null hypothesis of no local regulation for a given gene in any of the tissues under study (corresponding to a hypothesis in level 1 of the tree), we applied Simes’ combination rule to the gene x tissues p-values just described. We then tested the global null hypotheses of no local regulation in any tissue for all the genes in our study, applying the Benjamini Hochberg procedure^63^ to control the FDR at the 0.05 level. For those genes for which we were able to reject the null hypotheses of no local regulation, we examined the tissue-specific p-values, applying the Benjamini Bogomolov procedure that allows the identification of significant findings controlling for the initial selection^64^. Finally, the individual SNPs responsible for regulation of the gene in each tissue were identified, again using a selection-adjusted threshold as described in Bogomolov et al.^24^ An R package implementing this procedure is available at http://www.bioinformatics.org/treeqtl/.^65^

We compared the number of eGenes identified in each tissue using the above procedure with the results of GTEx (Analysis Release V6; dbGaP Accession phs000424.v6.p1). We downloaded all eQTL association results for tissues in common with our study, and applied this same hierarchical procedure to the GTEx results to identify eGenes.

#### Association between local eQTLs and genomic features

We estimated the possible enrichment of eQTLs in exons, introns, flanking regions, intergenic regions, and regulatory regions using logistic regression in a generalized linear mixed model (GLMM), using the GMMAT software^66^. We categorized each SNP in two binary dimensions (local eQTL and located in or near a given region). A SNP was considered a local eQTL if it was associated (at Bonferroni significance thresholds) to gene expression of a gene within 1 Mb, in any tissue, in either Dataset. Local eQTL status was the outcome variable, and a separate GLMM logistic regression performed for each region. A matrix of r^2^ values among all SNPs was included as a random effect to account for lack of independence among SNPs. GLMMs are computationally very demanding and the full set of 497,163 SNPs could not analyzed in one model. We LD pruned the SNP data, agnostic to eQTL status and region, and used 18,464 genome-wide SNPs based on LD pruning the entire set of 497,163 SNPs at r^2^<0.6 in 14 unrelated individuals. This SNP set included 1,202 local eQTLs.

#### Enrichment of local eQTLs in near TSS/TES

Our examination of potential enrichment of local eQTLs near TSS/TES gene regions was purely descriptive and involved no hypothesis testing. We restricted our summary to the 27,196 genes that were <0.5 Mb in size, and the 426,403 SNPs that were within 200kb of the TSS/TES of these genes (or in between the TSS/TES). In this set of 426,403 SNPs, 17,595 were local eQTLs to one (or more) of the 27,196 genes (at Bonferroni significance levels), in one (or more) tissues in either Dataset, and were within 200 Kb of the TSS/TES of the gene(s) to which they were associated. For each gene, we created 10 Kb distance bins on either side of the TSS/TES, and tallied the proportion of SNPs in the bin that were local eQTLs for the gene. As the distance between TSS and TES varied by gene, we binned distances in this area by deciles of the total distance.

## Data Availability

The RNA-Seq datasets generated in the current study are available in the NCBI Gene Expression Ominibus repository, https://www.ncbi.nlm.nih.gov/gds/?term=PRTNA219198. The other data sets, microarray and genotype, analysed during the current study are available in the NCBI Gene Expression Ominibus repository, https://www.ncbi.nlm.nih.gov/geo/query/acc.cgi?acc=GSE15301 (microarray data) and the EMBL-EBI, https://www.ebi.ac.uk/ena/data/view/ERP008917 (genotype data).

**Supplementary Informationis** linked to the online version of the paper at www.nature.com/nature.

## Acknowledgements

Thanks to Stephanie Groman for assistance with tissue resources. Thanks to Tara Chavanne, Kelsey Finnie, Margaret Long, Jean Gardin and Dianna Swaim for technical assistance with necropsies and tissue collections. This work was supported by the following grants, all from the U.S. National Institutes of Health: U54HG00307907 (to RKW); P40RR019963/ÜD010965 (to JRK); R01RR016300/ÜD010980 (to NBF); R37MH060233 (to Daniel Geschwind); UL1DE019580 (to Robert Bilder); PL1NS062410 (to Christopher Evans); RL1MH083270 (to JDJ); P30NS062691 (to NBF and GC); R01MH101782 (to CS and EE). RN, BLA, PF acknowledge support from Wellcome Trust PF, BLA, RN acknowledge support from the Wellcome Trust (grant number WT108749/Z/15/Z) and the European Molecular Biology Laboratory. ESW was supported by an EMBO Advanced Fellowship (aALTF1672-2014).

## Author contributions

AJJ, JRK, GMW, KD, RKW, JDJ, WW, RPW, and NBF designed the study. AJJ, IZ, OWC, JD, LAF, SF, AEF, YSH, VR, CAS, JDJ, GC, and RPW produced the data. AJJ, IZ, SKS, CP, RMC, EE, LAF, SF, YSH, VR, CAS, HS, DV, BLA, PF, RN, ESW, JB, TDD, MB, YB, CS, and GC analyzed the data. OWC, JD, and MJJ managed data and samples. AJJ, SKS, and NBF wrote the paper. All authors reviewed the final draft.

## Author Information

Reprints and permissions information is available at www.nature.com/reprints

The authors declare that they have no competing financial interests

Correspondence and requests for materials should be addressed to

